# Correlation of OCT-Based Radiomic Signatures With Dose-Associated Radiation Response in Tumor Spheroids

**DOI:** 10.64898/2026.07.08.737210

**Authors:** Mark Arndt, Rico Hansler, Luca Tirinato, Andrey Tkachenko, Joao Seco, Werner Nahm, Ute Schepers, Maria Francesca Spadea

## Abstract

**Background:** Three-dimensional tumor spheroids are an established radiobiology model, but scalable, reproducible readouts of dose-dependent radiation response are lacking. We evaluated whether optical coherence tomography (OCT) radiomics can quantify dose-associated response in spheroids, and how it compares with conventional brightfield morphology.

**Methods:** This in vitro, cross-sectional study used SAS oral squamous cell carcinoma spheroids seeded at two densities (5000 and 10000 cells), irradiated at 0 to 12 Gy, and imaged on days 1 to 11 post-irradiation. Each OCT acquisition yielded co-registered structural-intensity and speckle-variance volumes. Radiomic features (shape, first-order, texture) were extracted with Radiomics.jl, filtered for repeatability, correlation-pruned, and ensemble-ranked. Dose correlation was assessed by repeated 5-fold cross-validation across five regressors, comparing brightfield-only (BF), OCT-only, and combined OCT+BF feature sets with paired Wilcoxon tests.

**Results:** OCT-only models consistently outperformed the BF baseline (median ***R*^2^** 0.77 to 0.85 versus 0.61 to 0.69; ***p <* 0.001** for all regressors). Adding brightfield to OCT gave no consistent benefit, reaching significance only for Random Forest (***p* = 0.026**, power 0.62). A compact shared feature subset combined brightfield area dynamics with OCT texture, shape, and speckle-variance descriptors, all showing low repeat-scan variability relative to cohort variability.

**Conclusions:** OCT radiomics provides a sensitive, reproducible, label-free high-throughput readout of spheroid radiation dose response that outperforms the current brightfield-based approach, without requiring concurrent brightfield acquisition.

**Key points:**

- OCT radiomics quantifies dose-dependent radiation response in 3D tumor spheroids.
- OCT-only models outperformed brightfield imaging in radiation dose prediction.
- Selected radiomic features showed high repeatability across repeated acquisitions.
- Adding brightfield to OCT gave no consistent performance benefit.
- Label-free OCT enables reproducible, high-throughput spheroid radiation-response assessment.

## 1 Background

Spheroids provide a biologically relevant and scalable model system that bridges the gap between 2D cell cultures and *in vivo* tissue. Their three-dimensional architecture reproduces cell–cell interactions and diffusion-driven gradients (e.g., oxygen, pH, nutrients), which can substantially influence radiotherapy treatment response and make spheroids an established model in radiobiology (Dubessy et al. 2000). For radiobiology studies and therapy optimization, it is therefore essential to quantify dose-dependent spheroid response accurately and reproducibly across cohorts. In this context, high-throughput readouts are crucial to enable statistically robust dose– response experiments and to evaluate many conditions without prohibitive time and cost (Kunz-Schughart et al. 2004; Brüningk et al. 2020).

Common optical readouts face important trade-offs when applied to 3D microtissues. Brightfield imaging is fast and inexpensive, but it provides only a 2D projection view and cannot directly resolve internal 3D structure or depth-dependent changes (Hsieh et al. 2025). Fluorescence microscopy can add molecular specificity, yet it typically requires exogenous probes whose delivery and interpretation can be complicated in thick specimens; furthermore, repeated excitation and staining can perturb live samples (e.g., via phototoxicity), and many fluorescence-based assays are inherently end-point, limiting longitudinal measurements (Huang et al. 2019; Purschke et al 2010). While confocal microscopy enables optical sectioning, acquiring dense *z*-stacks is time-intensive and scanning-based imaging can be slow for high-throughput studies; in addition, scattering in thick samples limits effective imaging depth and leads to depth-dependent signal loss (Hsieh et al. 2025; Leary et al. 2018).

Optical coherence tomography (OCT) is emerging as a complementary imaging approach, providing label-free, depth-resolved volumetric imaging at micrometer-scale resolution, enabling non-destructive assessment of spheroid structure (Huang et al 1991; Fercher et al. 2003; Huang et al. 2019). Beyond structural information, temporal fluctuations in OCT intensity can provide contrast related to underlying tissue and cellular dynamics. Such signal variations can therefore serve as a source of quantitative imaging biomarkers, complementing purely morphological assessment (Abd El-Sadek et al. 2020, 2021, 2024). To capture this information in a systematic and reproducible way, quantitative feature extraction approaches are required. Radiation-induced biological alterations are known to manifest as spatially heterogeneous changes in the internal architecture of spheroids, including necrotic core expansion, cell death gradients, and alterations in microenvironmental dynamics, that evolve over days following irradiation and scale with delivered dose (Dubessy et al. 2000; Brüningk et al. 2020). Such changes are fundamentally three-dimensional and are expected to be largely invisible to brightfield projection imaging, which captures only a 2D outline of a complex volumetric process. OCT is therefore a promising candidate for resolving internal structural and dynamic changes associated with irradiation in a label-free, non-destructive manner. However, translating raw OCT volumes into reproducible, quantitative measures of dose-dependent spheroid response requires a systematic analytical framework.

Radiomics, originally established in radiological imaging, can bridge this gap by introducing a structured and reproducible pipeline that converts raw OCT volumes into quantitative, comparable descriptors. Applied to OCT, radiomics might have the potential to reveal subtle patterns and heterogeneity that may correlate with biological or treatment-related system behavior. By converting volumetric image data into high-dimensional descriptors of texture, shape, and intensity distribution, radiomics could enable the systematic decoding of subtle, dose-dependent patterns encoded in the OCT signal. The combination is therefore not incidental, as irradiation-induced changes modulate the OCT signal in ways that radiomics is designed to capture, and the dose-dependence of that signal provides a principled, experiment-defined target for quantitative validation.

Building on this rationale, the central research question of this study is to determine how quantitative radiomic descriptors derived from OCT, alone and in combination with complementary brightfield features, contribute to the assessment of dose-dependent spheroid response beyond conventional morphology-based readouts. We hypothesize that the combination of structural OCT information, dynamics-sensitive speckle-variance (SV) contrast, and brightfield morphology captures complementary aspects of spheroid response to irradiation and therefore enables a more sensitive and comprehensive characterization of that dose-dependent response. The novelty of this work lies in integrating OCT-based radiomics with radiation-response assessment in a label-free multimodal framework, with the aim of uncovering imaging biomarkers that provide deeper insight into treatment-induced alterations and improve the assessment of dose-dependent spheroid response beyond BF-only analysis. To investigate this hypothesis, we developed a multimodal analysis pipeline that combines radiomic features, derived from intensity and SV OCT images, with brightfield measurements of tumor spheroids. The added value of this multimodal approach was assessed in comparison with conventional BF-only analysis.

## 2 Methods

### Study design

This was an in vitro, cross-sectional imaging study of SAS human tongue squamous cell carcinoma tumor spheroids. As the study used an established human-derived cell line and did not involve human participants, identifiable human material, or animals, ethics committee approval and informed consent were not applicable. An overview of the full experimental and analysis pipeline is provided in Fig. 1.

**Fig. 1:**
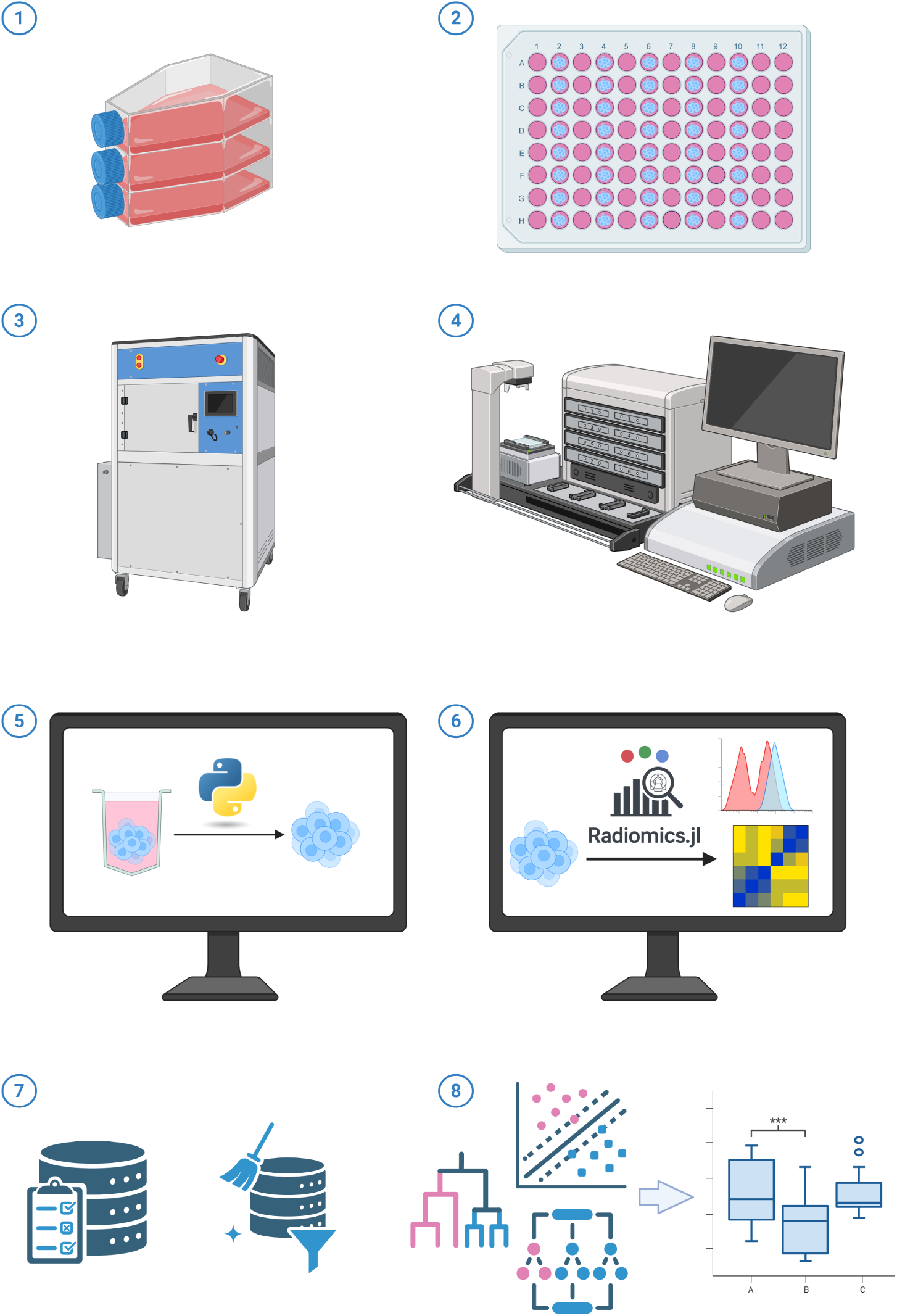
Overview of the experimental and analysis pipeline. (1) SAS oral squamous cell carcinoma cells are expanded in stan6dard monolayer culture. (2) Spheroids are formed by seeding 5000 or 10000 cells per well in 96-well agarose plates. (3) Plates are irradiated with doses of 0 to 12 Gy using a cabinet X-ray irradiator. (4) At each post-irradiation time point, spheroids are imaged by both brightfield microscopy and OCT; for each lateral position, repeated B-scan frames are acquired to directly compute speckle-variance maps alongside the structural intensity volumes. (5) OCT volumes and SV maps are segmented and processed using a custom Python pipeline. (6) Radiomic features (shape, first-order, and texture families) are extracted from segmented intensity volumes and variance maps using Radiomics.jl. (7) Features undergo robustness filtration, correlation pruning and ensemble ranking to identify a compact, non-redundant predictor set. (8) Dose correlation performance is evaluated via repeated 5-fold cross-validation across multiple regression model families, with three-way statistical comparison of BF-only, OCT-only, and combined feature sets.

**Fig. 2:**
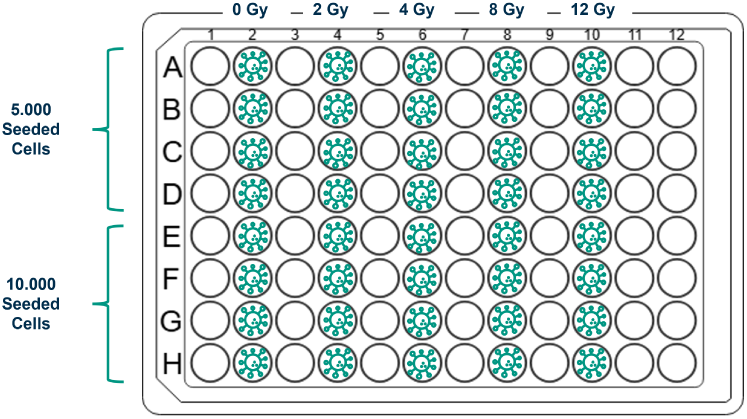
Plate layout: 5000-cell spheroids (A–D) and 10000-cell spheroids (E–H) irradiated at 0, 2, 4, 8, and 12 Gy (4 replicates/condition).

### 2.1 Cell culture

The cells, SAS human tongue squamous cell carcinoma, were cultured in Dulbecco’s Modified Eagle Medium (DMEM) containing 10% fetal bovine serum (FBS) and 1% penicillin/streptomycin at 37 *^◦^*C and 5% CO_2_. Passages 17 to 19 of the cells were used for the experiments. Cell cultures were routinely screened for mycoplasma contamination and confirmed to be negative before being used in the experiments.

### 2.2 Spheroid assay

Cells were harvested at 90% confluency for use in generating spheroids. Spheroids were formed by seeding 5000 or 10000 cells per well in 100 µl of cell culture medium in a 96-well agarose plate (Onozato et al. 2017). The plates were prepared using 50 µl of 1.5% agarose solution per well. After seeding, the spheroids were maintained for an additional 6 days to reach the target size.

To facilitate a longitudinal assessment, repeated measurements of the same spheroids would be preferable. However, in our current workflow, OCT imaging requires sample handling that could increase contamination risk. We therefore adopted a cross-sectional design: independent spheroid plates were prepared for each measurement day under identical seeding and culture conditions, ensuring comparable initia conditions across timepoints while avoiding repeated handling of the same samples.

### 2.3 Irradiation

Spheroids were imaged by brightfield immediately before irradiation (baseline) and then irradiated with doses of 0, 2, 4, 8, or 12 Gy. Follow-up imaging was performed at 1, 4, 6, 8, and 11 days post-irradiation. At days 1, 4, and 6, brightfield imaging only was performed. At days 8 and 11, both brightfield and OCT imaging were performed; these time points were designated as the primary model-training time points. Day 8 used five plates and day 11 used four plates, corresponding to *n* = 20 and *n* = 16 spheroids per condition, respectively (each plate contributing four replicates per condition).

Irradiations were performed using a cabinet-based photon (X-ray) irradiator (Faxitron MultiRad 225 (Faxitron 2024)), operated at a dose rate of approximately 2 Gy min*^−^*^1^ at the sample position.

### 2.4 Brightfield

Brightfield images were acquired to obtain 2D morphological readouts (spheroid diameter and projected area) as complementary predictors alongside OCT radiomics. Images were recorded at 10*×* magnification. To avoid inaccuracies from fully automated contour detection in brightfield (e.g., due to low contrast, irregular borders, or debris) and to best replicate the manual evaluation procedure, spheroid diameter and projected area were measured manually in ImageJ (Schneider et al. 2012). For post-irradiation days, spheroid diameter and projected area were measured and, together with their absolute and relative changes compared to the pre-irradiation baseline, used as features.

To statistically assess whether brightfield size metrics captured a dose-dependent effect, relative area and diameter change (relative to the pre-irradiation baseline) were compared across the five dose groups (0, 2, 4, 8, and 12 Gy) at each post-irradiation imaging day using the non-parametric Kruskal–Wallis test, performed separately for the pooled cohort and for each seeding-density subgroup (5000-cell and 10000-cell spheroids). For days at which the omnibus test reached significance (*p <* 0.05), post-hoc pairwise Mann–Whitney U tests comparing each irradiated dose group against the 0 Gy control were performed.

### 2.5 Optical coherence tomography

Optical coherence tomography (OCT) is a label-free, non-destructive imaging technique that provides depth-resolved, micrometer-scale structural information in optically scattering samples (Huang et al. 1991; Fercher et al. 2003). OCT is based on low-coherence interferometry: backscattered light from different depths in the sample interferes with light from a reference arm only when the optical path lengths match within the coherence length, enabling ranging along the axial direction. A single depth scan is referred to as an A-scan; lateral scanning yields a cross-sectional B-scan, and dense raster scanning enables volumetric (3D) imaging (Huang et al. 1991; Fercher et al. 2003).

Temporal fluctuations of the OCT intensity can be exploited to obtain dynamics- sensitive contrast. In this work, we use *speckle variance* (SV), a variance-based metric widely used in dynamic OCT/OCT angiography to visualize and quantify tempora signal fluctuations arising from microscopic motion within the sample (Mariampilla et al. 2008, 2010; Gorczynska et al. 2016). For each voxel/pixel, let *I_k_* denote the OCT intensity in repeat *k*. Speckle variance is computed as the temporal variance of *I_k_*:

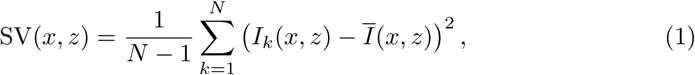

where 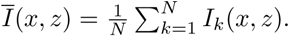 In this study, *N* = 20 repeated B-scan frames were acquired at each lateral position to compute SV. Each OCT acquisition therefore yields two co-registered volumetric outputs, a structural intensity volume encoding depth-resolved backscatter, and a speckle-variance volume encoding temporal intensity fluctuations. A three-dimensional rendering of a representative SV volume at the highest dose condition (12 Gy), with a wedge cut revealing the internal variance distribution, is shown in Fig. 3.

**Fig. 3:**
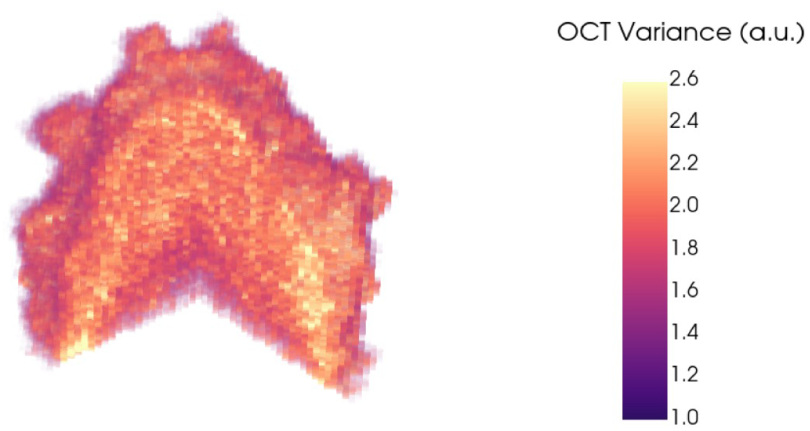
Three-dimensional volume rendering of the OCT speckle-variance (SV) map for a representative spheroid irradiated at 12 Gy. A wedge cut has been applied to expose the internal variance distribution. Color encodes local SV intensity, with higher values indicating regions of stronger temporal fluctuation and lower values indicating more static regions. The spheroid diameter is approximately 400 µm.

Regions exhibiting stronger temporal fluctuations yield higher SV values. SV contrast depends on acquisition parameters such as the number of repeats *N* and the inter-frame interval; bulk motion can inflate variance and reduce specificity, motivating stable sample mounting and (when needed) motion correction for quantitative use (Mariampillai et al. 2010; Gorczynska et al. 2016). We hypothesize that these dynamics-sensitive fluctuations reflect treatment-induced changes in spheroid viability and microenvironmental activity, thereby providing complementary information to structural OCT. As such, SV may help reveal subtle irradiation-induced effects and improve imaging-based assessment of dose-dependent spheroid response when integrated with structural and morphological descriptors.

For spheroid imaging, OCT data were acquired using a spectral-domain OCT system (Thorlabs Ganymede series; item GAN612C1/M (Thorlabs 2025)) with a center wavelength of 880 nm, operated at a 5 kHz A-scan rate, yielding a B-scan acquisition rate of 33 Hz. Volumes were acquired with a voxel spacing of 6.0 µm laterally and 3.5 µm axially. Axial scan depth was corrected for the refractive index of water. During imaging, spheroids were removed from the incubator and positioned and oriented in a standardized manner using a custom temperature-controlled sample holder/stage maintained at around 37 *^◦^*C, ensuring consistent imaging geometry across acquisitions. The acquisition time per spheroid was approximately 1 to 2 min. To compute speckle-variance maps, *N* = 20 repeated B-scan frames were acquired at each latera position (Eq. 1), and the temporal fluctuations of the OCT intensity were used to form variance-derived contrast (Gorczynska et al. 2016).

### 2.6 Image processing: segmentation

OCT volumes were standardized using a cropping and masking workflow. First, each spheroid stack was inspected in a custom cropping interface that displays XZ slices and allows manual bounding-box selection with adjustable Y-limits to control the longitudinal extent; the selected crop bounds were saved per well identifier. These bounds were then applied during TIFF-to-NIfTI conversion to enforce consistent fields-of-view and voxel geometry across samples, with voxel spacing taken from image metadata when available and preserved in the NIfTI affine.

Second, spheroid masks were generated using Otsu thresholding (Otsu 1979) on the OCT intensity, followed by 3D morphological closing/opening, removal of small components, and retention of the largest connected component to suppress noise and spurious structures. Additional z-slice trimming was applied to remove low-signal regions near the volume boundaries. Each mask was visually reviewed and either saved or discarded, yielding a curated segmentation set. Accepted masks were saved as NIfTI and used to restrict radiomics to spheroid tissue, reducing background and boundary artifacts.

### 2.7 Image processing: radiomics

#### Radiomics and feature matrix construction

Radiomic features were extracted from OCT intensity volumes and SV maps using Radiomics.jl (Zaffino et al. 2026) then merged per sample and timepoint. The following feature classes were enabled shape (only for Intensity), first-order, GLCM, GLRLM, GLSZM, and GLDM. Shape features were computed only once from the Intensity volumes because the same mask is used for the variance-derived data, making shape features redundant for those volumes.

To assess feature robustness against repeated acquisition and pipeline variability, a dedicated repeatability experiment was performed in which a single spheroid was imaged six times under unchanged conditions. Features were extracted for each repeat using the identical preprocessing and extraction pipeline. For each feature, the IQR/Median across the six repeats was computed. In addition, the within-spheroid variability was summarized by the feature IQR across repeats and compared to the population-level IQR of the same feature across all spheroids, providing a relative variability context (i.e., whether repeat-scan variability was small compared to biological/condition variability).

The brightfield measurements (diameter, area, and temporal derivatives) were matched by sample identifier across days. The final feature matrix combined radiomics and brightfield predictors. For feature selection, the analysis focused on the final two days of imaging (day 8 and day 11) to reduce temporal confounding.

It is important to note that brightfield features include both absolute measurements and change metrics relative to the pre-irradiation baseline, providing an implicit within-spheroid temporal reference. OCT features, by contrast, are derived from a single volumetric acquisition per timepoint with no equivalent baseline scan, meaning OCT operates without this longitudinal anchor. This asymmetry should be considered when interpreting the relative performance of the two modalities.

#### Radiomic OCT feature robustness filtration

To assess OCT feature robustness, a single spheroid was imaged six consecutive times under identical conditions, and features were extracted using the same processing pipeline. For each feature, repeatability was quantified by the absolute interquartile range (IQR) and the relative variability (IQR*/*median) across the six repeated acquisitions. Features with an IQR*/*median exceeding 35% were excluded, as they were considered insufficiently robust and likely dominated by measurement noise. Applying this criterion led to the removal of 24 out of 198 extracted features.

Missing values were imputed with median statistics and only numeric predictors were retained. To enable a fair comparison between OCT-only and combined OCT+BF models, all subsequent feature-selection steps, correlation pruning, univariate screening, ensemble ranking, and the Top-*N* sweep, were performed independently for two separate feature-set definitions: (i) an OCT-only feature pool, containing exclusively OCT-derived radiomic features, and (ii) a combined OCT+BF feature pool. This separation ensures that an OCT feature is not excluded from the OCT-only analysis merely because it was pruned in favor of a correlated brightfield feature during the combined-pipeline selection; the OCT-only feature set is therefore derived from its own independent selection process rather than being a post-hoc subset of the combined selection. Within each feature-set definition, multicollinearity was reduced by pruning features with Spearman correlation magnitude *|ρ| ≥* 0.95, keeping the higher-variance feature from each correlated pair.

Regression models evaluated were Random Forest (RF) and Extra Trees (ET) (Breiman 2001; Geurts et al. 2006), Support Vector Regression (SVR; RBF kernel) (Smola and Schölkopf 2004), and gradient-boosted decision trees (LightGBM and XGBoost) (Ke et al. 2017; Chen and Guestrin 2016). All models were assessed using repeated 5-fold cross-validation, each with 5 repeats, to ensure the stability of the performance results (Kohavi 1995).

Within each feature-set definition, univariate screening quantified feature–dose associations using Spearman correlation, mutual information, and F-test scores Model-based importance was assessed using cross-validated permutation importance computed independently for each of the five regression models above. For each model, permutation importance scores were min-max normalized to the [0, 1] range across features, and the normalized scores were then aggregated across models into a single model-based importance score. This aggregated model-based importance score contributed 85% of the weight of the overall ensemble feature-ranking score, with the normalized Spearman correlation, mutual information, and F-test scores each contributing 5% (15% in total) of the remaining weight.

Rather than fixing a single feature count, a Top-*N* sweep was performed independently within each feature-set definition, and the best *N* was selected *per model and per feature-set definition* using the median cross-validated *R*^2^ (with IQR). This approach was preferred over a fixed feature count to account for the fact that different models, and different feature-set definitions, may require different levels of predictor complexity to generalize well. For the combined OCT+BF feature set, the optimal values were *N*_RF_ = 20, *N*_ET_ = 25, *N*_SVR_ = 7, and *N*_GBM_ = 20 (both gradient-boosted models). For the independently selected OCT-only feature set, the optimal values were *N*_RF_ = 15, with *N* = 20 for Extra Trees, SVR, and both gradient-boosted models.

To mitigate overfitting, regularization strategies were applied during model configuration. For tree ensembles, complexity was constrained by limiting depth and enforcing minimum sample requirements for splits/leaves (e.g., max depth, min samples split, min samples leaf), and by using feature subsampling per split (e.g., max features=’sqrt’) (Breiman 2001; Geurts et al. 2006). For gradient boosting, regularization was achieved via shrinkage (learning rate), tree-depth constraints, and feature subsampling (Ke et al. 2017; Chen and Guestrin 2016).

A three-way feature-set comparison was performed: (i) combined OCT+BF features, using the model-specific Top-*N* subset selected from the combined feature-set definition; (ii) OCT-only features, using the model-specific Top-*N* subset selected independently from the OCT-only feature-set definition; and (iii) BF-only features (baseline). Performance differences were assessed with paired Wilcoxon tests on per-fold *R*^2^ values (Wilcoxon 1945), complemented by bootstrap resampling for uncertainty estimation and power assessment (Efron and Tibshirani 1993).

## 3 Results

A total of 130 spheroids were included in the day-8 analysis and 142 spheroids in the day-11 analysis, distributed across the five irradiation doses (0, 2, 4, 8, and 12 Gy) and the two seeding densities (5000 and 10000 cells per well). Brightfield imaging was performed at all six imaging time points (baseline and days 1, 4, 6, 8, and 11), whereas OCT imaging was restricted to days 8 and 11, the model-training time points.

### 3.1 Brightfield results

Spheroid projected diameter and area were measured in brightfield before irradiation (day 0) and on post-irradiation days 1, 4, 6, 8, and 11. For each dose–seeding condition, measurements across replicate spheroids were summarized by the median and interquartile range (IQR), and size dynamics were visualized as the relative change with respect to the pre-irradiation baseline (Figs. 4 and 5). A clear seeding effect was observed: small spheroids (5000 cells) expanded rapidly already by day 1 and generally showed larger early increases, whereas large spheroids (10000 cells) exhibited slower early dynamics and, in several irradiated groups, an initial compaction (negative change) before later regrowth.

**Fig. 4:**
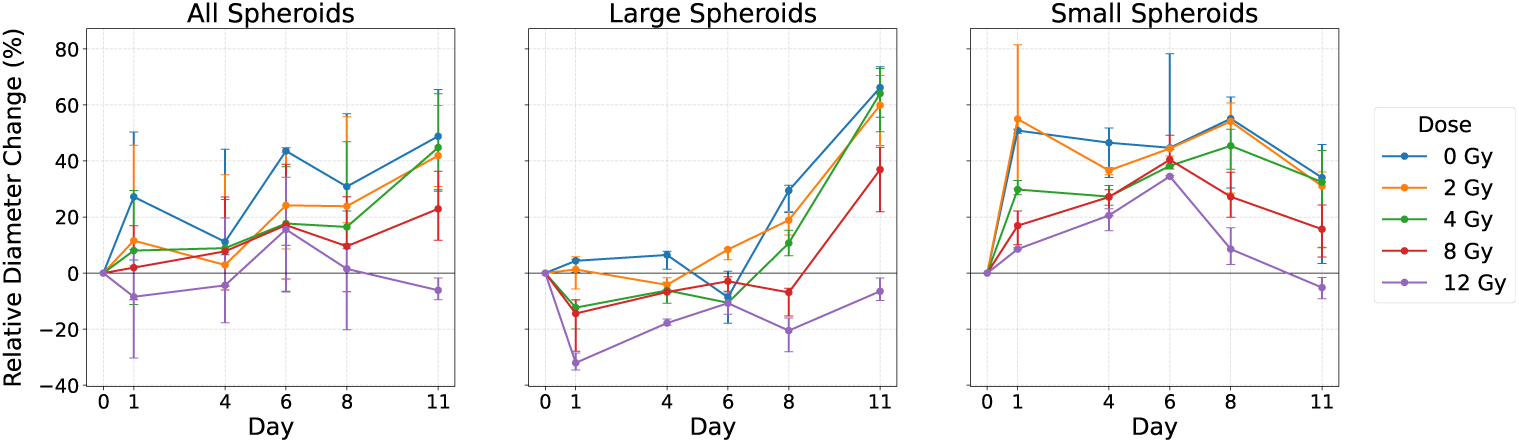
Relative diameter change (%) compared to the pre-irradiation day (median *±* IQR).

**Fig. 5:**
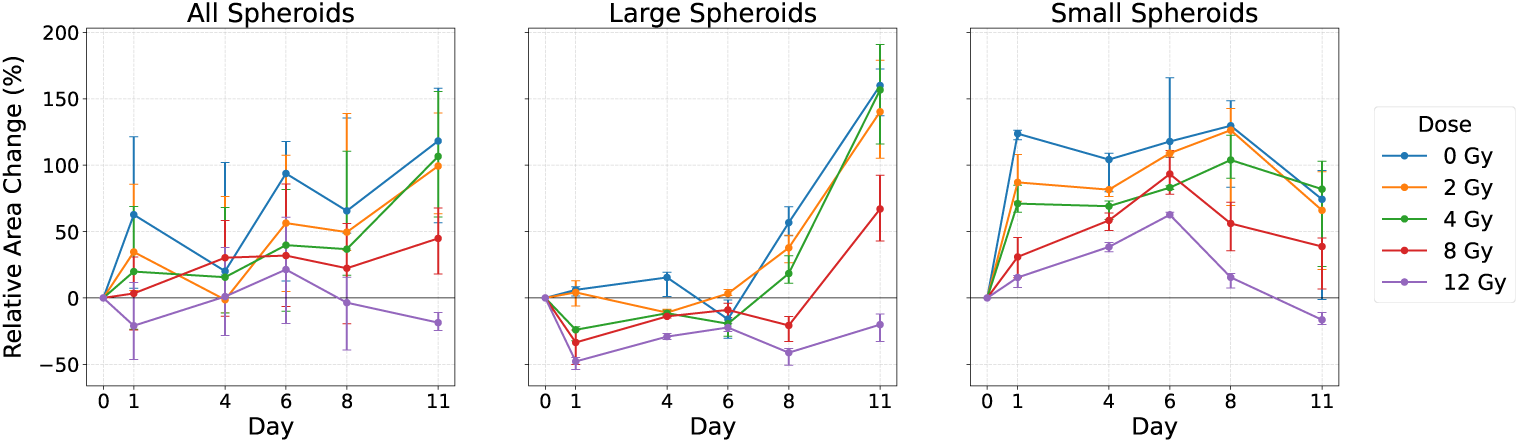
Relative area change (%) compared to the pre-irradiation day (median *±* IQR)

Across both seeding densities, irradiation suppressed growth in a dose-dependent manner. Lower doses (0 to 4 Gy) retained stronger expansion, while 8 Gy showed reduced growth and 12 Gy produced the strongest inhibition and, in some cases net shrinkage at late time points (day 11). In addition to these size changes, late time points (days 8 to 11) showed qualitative boundary alterations in some irradiated spheroids, with fringes/irregular halos at the rim that were less prominent in controls

These trends were confirmed statistically using Kruskal–Wallis tests across dose groups at each imaging day, performed on the relative area and diameter change with respect to the pre-irradiation baseline, i.e., the same baseline-normalized metrics used as brightfield predictors elsewhere in this study. At days 0, 1, 4, and 6, the omnibus test showed no consistent dose effect on either metric, in any cohort grouping (*p >* 0.07). From day 8 onward, the omnibus test was highly significant for both metrics across all cohort groupings (*p <* 0.001). Post-hoc Mann–Whitney U tests against the 0 Gy control showed that this separation was driven by the higher dose groups: 8 and 12 Gy differed significantly from control at both day 8 and day 11 (*p <* 0.001, pooled cohort), whereas 2 Gy never reached significance and 4 Gy reached significance only in the large-spheroid subgroup at day 8. Notably, since these metrics already incorporate a within-spheroid baseline correction, the absence of a detectable dose effect below 4 Gy at any timepoint cannot be attributed to a missing temporal reference, but instead reflects a genuine limit of brightfield size readouts to resolve dose differences in this range.

This pattern, namely the inability of brightfield size metrics to resolve a dose-dependent effect up to 4 Gy (and, in the smaller-spheroid subgroup, up to 8 Gy) at any timepoint, together with the consistent and strong separation that emerges only from day 8 onward, motivated restricting the subsequent radiomics dose-correlation analysis to days 8 and 11, the earliest timepoints at which a measurable brightfield effect became reliably detectable.

### 3.2 OCT-derived radiomic results

#### Representative OCT en face projections across dose

To visually illustrate dose-associated changes captured by OCT beyond the single 12 Gy example shown in Fig. 3, Fig. 6 shows representative en face projections (mean intensity and mean speckle variance, each averaged over the depth (*z*) axis) of individual spheroids at day 11 across the five irradiation doses (0, 2, 4, 8, and 12 Gy), normalized on a shared scale to allow direct visual comparison across doses. Consistent with the boundary alterations described qualitatively in the brightfield results above, spheroid outlines become visibly more irregular and fringed with increasing dose in both the intensity and variance projections, most apparent at 8 and 12 Gy. Beyond this boundary effect, the spatial pattern of the SV signal also changes: at 0 to 4 Gy, the en face variance maps show a smooth, broadly concentric pattern of high variance across the central region with a thin low-variance rim, whereas at 8 and 12 Gy this pattern becomes visibly more heterogeneous, with patchy, irregularly distributed regions of elevated variance rather than a single smooth interior plateau. This qualitative shift toward greater spatial heterogeneity at higher doses is consistent with the prominence of gray-level non-uniformity texture features, derived from both the intensity and variance volumes, among the top-ranked predictors identified by the ensemble feature ranking (see below).

**Fig. 6:**
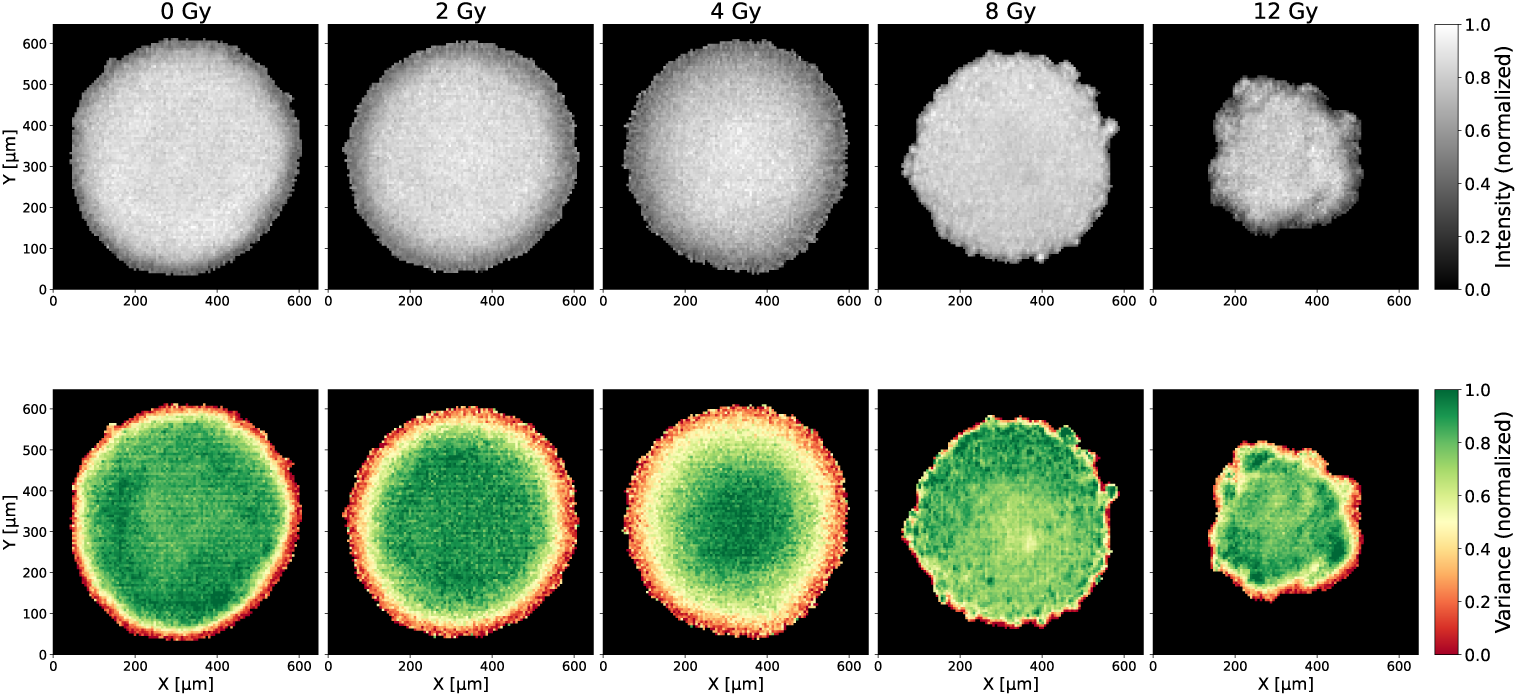
Representative en face OCT projections of individual spheroids at day 11, at each irradiation dose, averaged over depth (*z*) and normalized on a shared scale across doses. Top row: mean intensity. Bottom row: mean speckle variance (SV). Spheroid boundaries become progressively more irregular with increasing dose, and the internal SV pattern shifts from a smooth, concentric distribution at low doses to a more spatially heterogeneous, patchy distribution at 8 and 12 Gy.

#### Cohort overview and feature ranking

The ensemble ranking for the combined OCT+BF feature set consistently placed brightfield-derived size metrics among the strongest predictors. To assess agreement across model families, the top-10 ranked features from this combined ranking were compared pairwise between all five models (Fig. 7). Pairwise overlap ranged from 5 to 9 shared features (out of 10) and reflected underlying model similarity: the two tree ensembles, RF and ET, shared 9 of their top-10 features, and the two gradient-boosted models, XGBoost and LightGBM, likewise shared 9. SVR shared comparably high overlap with ET (9 features) but markedly lower overlap with both gradient-boosted models (5 to 6 features), indicating that its ranking diverges most strongly from the boosting-based models rather than from all other model families uniformly.

**Fig. 7:**
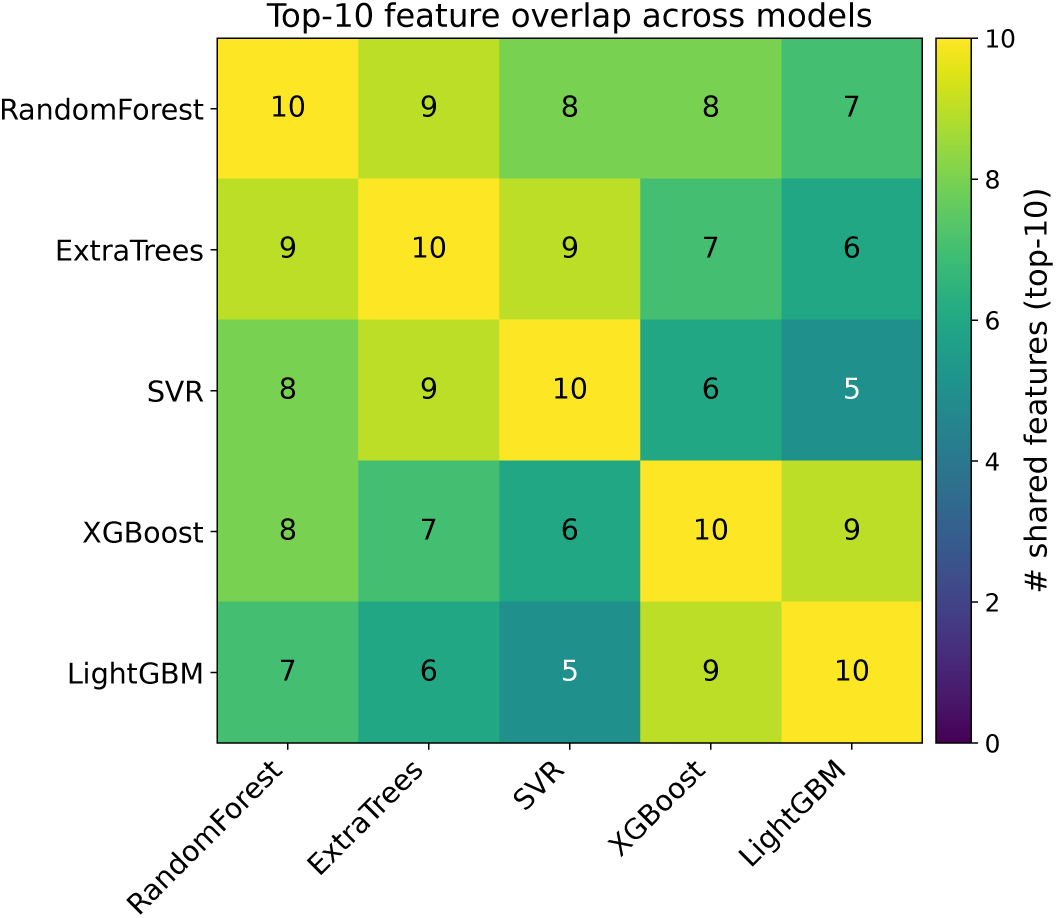
Pairwise overlap in the top-10 ranked features (out of 10) across the five regression models used for feature selection.

Five features were included in the top-10 of every model: Brightfield Area, Intensity original glrlm GrayLevelNonUniformity, Intensity original shape Sphericity, Variance original firstorder Skewness, and Variance original gldm GrayLevelNonUniformity. These are the same four OCT-derived descriptors examined in the repeatability analysis below (Table 1).

**Table 1:** Repeatability and variability context for selected radiomic features. Repeat scans: one spheroid imaged *N* = 6 times.

| Feature | IQR (N=6) | IQR/Median (N=6) | IQR (cohort) | $\Delta$ IQR (%) |
| --- | --- | --- | --- | --- |
| `Intensity_original_glrlm_GrayLevelNonUniformity` | 310 | 0.047 | 76000 | +25000 |
| `Intensity_original_shape_Sphericity` | 0.012 | 0.044 | 0.090 | +670 |
| `Variance_original_firstorder_Skewness` | 0.0065 | 0.036 | 0.26 | +3900 |
| `Variance_original_gldm_GrayLevelNonUniformity` | 1400 | 0.032 | 510000 | +37000 |

#### Feature robustness

To quantify repeatability of key radiomic features identified by the combined OCT+BF ranking, one spheroid was imaged six times under unchanged conditions and processed with the identical extraction pipeline. For each feature, the relative dispersion across repeats was summarized as the IQR-to-median ratio (IQR*/*median), alongside the absolute IQR across the six acquisitions (Table 1). Repeat-scan IQRs were additionally compared to the population-level IQR of the same features across the full cohort (Table 1).

Across the four evaluated features, relative dispersion on repeat scans was low and comparable for all of them (IQR*/*median = 0.032 to 0.047). Intensity original glrlm GrayLevelNonUniformity showed the highest relative dispersion among thefour (0.047), and Variance original gldm GrayLevelNonUniformity the lowest (0.032), with Intensity original shape Sphericity (0.044) and Variance original firstorder Skewness (0.036) falling in between. This indicates that, despite being texture- and intensity-distribution-sensitive descriptors, all four selected features were extracted with comparably low variability under unchanged acquisition conditions.

Cohort-level variability substantially exceeded repeat-scan variability for all four features, though to differing degrees. The increase was most pronounced for the two gray-level non-uniformity descriptors, Variance original gldm GrayLevelNonUniformity (ΔIQR *≈* +37000%) and Intensity original glrlm GrayLevelNonUniformity (*≈* +25000%), and comparatively smaller, though still substantial, for Variance original firstorder Skewness (*≈* +3900%) and Intensity original shape Sphericity (*≈* +670%). This pattern indicates that between-spheroid, dose-related variability dominates over repeat-scan measurement variability for all four features, supporting their use as reproducibly extracted predictors of dose-dependent spheroid response.

### Dose correlation performance

All models outperformed the BF-only baseline (Table 2). With OCT-only feature selection performed independently from the combined OCT+BF selection (see Methods), OCT-only models achieved median *R*^2^ values ranging from 0.77 to 0.85 across regressors, compared to 0.61 to 0.69 for BF-only models, with statistically significant improvements in all OCT-only versus BF comparisons (*p <* 0.001; Table 2). Models combining OCT and brightfield (OCT+BF) achieved median *R*^2^ values ranging from 0.80 to 0.84. The OCT+BF versus OCT-only difference reached significance only for Random Forest (*p* = 0.026), with comparatively low power (0.62), and was not significant for the remaining four models (Extra Trees, *p* = 0.055; SVR, *p* = 0.770; XGBoost, *p* = 0.670; LightGBM, *p* = 0.280 Table 2). The median *R*^2^ values for OCT-only and OCT+BF were correspondingly close for these four models, with LightGBM showing a marginally higher OCT-only median (0.85 versus 0.84) and Extra Trees, SVR, and XGBoost showing a marginally higher OCT+BF median; given the absence of statistical significance, none of these differences are interpreted as practically meaningful.

**Table 2:** Best feature-set size (*N*, selected independently for the OCT-only and combined OCT+BF feature sets), and *R*^2^, MAE, and RMSE performance (median (IQR)) for brightfield-only (BF), OCT-only, and combined OCT+BF models, with statistical tests comparing (i) OCT-only versus BF-only and (ii) OCT+BF versus OCT-only.

|  | RF | ET | SVR (RBF) | LightGBM | XGBoost |
| --- | --- | --- | --- | --- | --- |
| Best $N$ (OCT+BF) | 20 | 25 | 7 | 20 | 20 |
| Best $N$ (OCT-only) | 15 | 20 | 20 | 20 | 20 |
| $R^2$ (BF-only) | 0.68 (0.60–0.76) | 0.66 (0.60–0.71) | 0.69 (0.64–0.77) | 0.65 (0.58–0.75) | 0.61 (0.59–0.73) |
| MAE (BF-only) | 1.71 (1.55–1.89) | 2.07 (1.86–2.16) | 1.63 (1.50–1.83) | 1.73 (1.48–1.95) | 1.74 (1.55–1.97) |
| RMSE (BF-only) | 2.29 (2.10–2.73) | 2.49 (2.29–2.69) | 2.30 (2.01–2.65) | 2.34 (2.04–2.78) | 2.44 (2.17–2.80) |
| $R^2$ (OCT-only) | 0.77 (0.75–0.83) | 0.78 (0.75–0.83) | 0.84 (0.81–0.86) | 0.85 (0.80–0.86) | 0.83 (0.82–0.87) |
| MAE (OCT-only) | 1.48 (1.39–1.59) | 1.49 (1.38–1.60) | 1.32 (1.19–1.35) | 1.30 (1.21–1.44) | 1.29 (1.21–1.38) |
| RMSE (OCT-only) | 1.95 (1.78–2.08) | 1.91 (1.81–2.01) | 1.72 (1.60–1.85) | 1.70 (1.60–1.85) | 1.74 (1.55–1.84) |
| $R^2$ (OCT+BF) | 0.80 (0.76–0.85) | 0.81 (0.76–0.84) | 0.84 (0.81–0.87) | 0.84 (0.80–0.86) | 0.83 (0.80–0.87) |
| MAE (OCT+BF) | 1.44 (1.38–1.53) | 1.48 (1.40–1.56) | 1.28 (1.25–1.41) | 1.25 (1.20–1.37) | 1.29 (1.16–1.36) |
| RMSE (OCT+BF) | 1.83 (1.75–1.96) | 1.87 (1.77–1.96) | 1.69 (1.57–1.79) | 1.74 (1.54–1.84) | 1.68 (1.57–1.90) |
| $p$ (OCT-only vs. BF) | $1 \times 10^{-5}$ | $1.8 \times 10^{-7}$ | $1.2 \times 10^{-7}$ | $6.0 \times 10^{-7}$ | $1.2 \times 10^{-7}$ |
| Power (OCT-only vs. BF) | 1.000 | 1.000 | 1.000 | 1.000 | 1.000 |
| $p$ (OCT+BF vs. OCT) | 0.026 | 0.055 | 0.770 | 0.280 | 0.670 |
| Power (OCT+BF vs. OCT) | 0.619 | 0.490 | 0.063 | 0.208 | 0.077 |

Figure 8 shows fold-wise *R*^2^ distributions for LightGBM, comparing the combined OCT+BF Top-20 feature set, OCT-only, and BF-only. Both OCT-only and OCT+BF yielded higher fold-wise *R*^2^ than BF-only, with a paired Wilcoxon test indicating a significant difference versus the BF-only baseline for OCT-only (*p <* 0.001; Table 2). For LightGBM, the OCT+BF versus OCT-only comparison was not significant (*p* = 0.280 Table 2), with OCT-only achieving a numerically higher median *R*^2^ (0.85 versus 0.84) Figure 8b shows predicted versus true irradiation dose for the combined OCT+BF Top-20 LightGBM model, pooled across cross-validation folds.

**Fig. 8:**
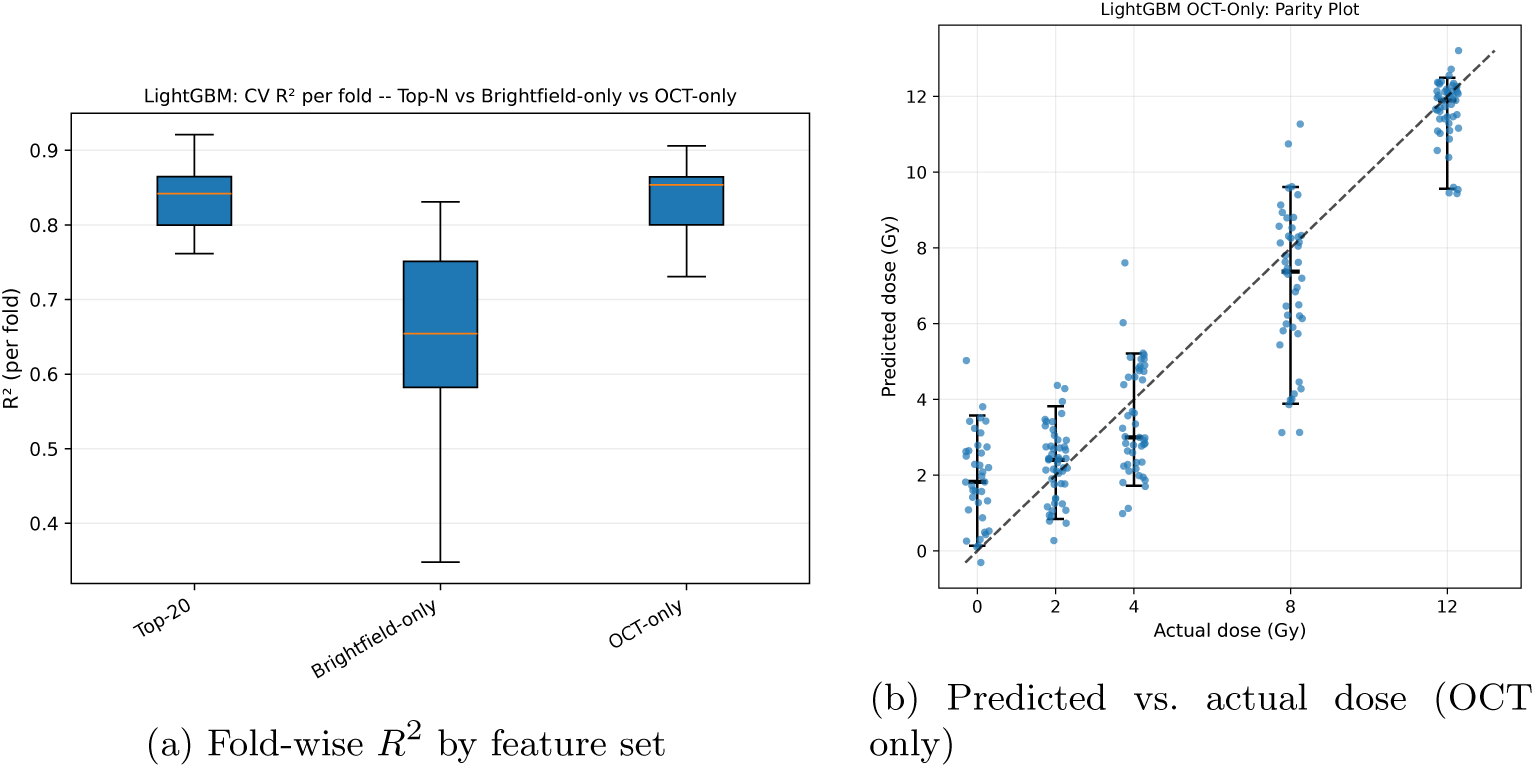
LightGBM cross-validated dose-correlation performance. (a) Fold-wise *R*^2^ for the combined OCT+BF Top-20 feature set, BF-only, and OCT-only. Boxes indicate the interquartile range with median (center line); whiskers denote the spread across folds. (b) Predicted versus true irradiation dose for the OCT-only feature set, pooled across cross-validation folds; the dashed line indicates perfect prediction. Reported *p*-values correspond to paired Wilcoxon tests versus the BF-only baseline.

Overall, the added value of combining brightfield with OCT radiomics, relative to OCT-only, was limited and largely model-dependent. The OCT+BF versus OCT- only difference reached statistical significance only for Random Forest, and even there with comparatively low power (0.62); for Extra Trees, SVR, XGBoost, and Light- GBM, the difference was not statistically significant (Table 2). For SVR and XGBoost OCT-only models additionally showed a narrower cross-validation IQR of *R*^2^ than the corresponding OCT+BF models, and LightGBM achieved a nominally higher median *R*^2^ with OCT-only than with OCT+BF. These results indicate that OCT radiomics alone is capable of matching, and in several cases equaling or marginally exceeding the performance of the combined OCT+BF pipeline. This supports OCT radiomics as a robust, self-sufficient, high-throughput readout of dose-dependent spheroid response that does not require concurrent brightfield acquisition.

## 4 Discussion

This study evaluated whether OCT-derived radiomics (from intensity volumes and SV maps) can serve as a quantitative, high-throughput readout of dose-dependent spheroid response in SAS spheroids, and how these descriptors compare to conventional brightfield size metrics. Across all evaluated model families, OCT-only features consistently outperformed the BF-only baseline in dose correlation (Table 2), supporting the premise that depth-resolved OCT encodes treatment-related information beyond 2D projected morphology.

The consistent superiority of OCT-only models over the brightfield baseline is noteworthy given that the two feature sets are not informationally equivalent. Brightfield change metrics encode a within-spheroid delta relative to the pre-irradiation size, which directly captures dose-dependent growth suppression and provides a strong longitudinal signal. OCT features, derived from a single snapshot per timepoint without an equivalent baseline volume, lack this explicit temporal reference. That OCT still outperforms brightfield under these conditions suggests that the structural and dynamic information encoded in a single OCT acquisition carries substantial dose-related signal, and that adding a pre-irradiation OCT baseline scan would likely improve performance further. This also motivates the prioritization of longitudinal OCT acquisition as a key direction for future work.

Brightfield measurements captured clear macroscopic trends: seeding density strongly influenced early growth kinetics, and irradiation suppressed expansion in a dose-dependent manner (Figs. 4 and 5). However, the BF-only baseline remained limited by its projection nature and sensitivity to boundary artifacts, which became more apparent at later time points where fringes and irregular halos were observed in some irradiated spheroids. Such border changes can bias diameter/area readouts and highlight the practical limitation of using 2D outline-based metrics as a standalone surrogate for 3D tissue response.

Feature ranking and cross-model feature overlap further indicate that the dose signal is distributed across complementary descriptors. The fact that all models selected the same small set of predictors combining brightfield size dynamics with OCT texture, shape, and SV-derived distribution features suggests that both macroscopic growth inhibition and structural/dynamic changes across the spheroid volume contribute to dose separability. In particular, sphericity quantifies the surface roughness of the segmented spheroid volume, with a lower value reflecting a more irregular, less smooth boundary. This is consistent with the qualitative boundary alterations described above for brightfield imaging, where late post-irradiation time points showed fringes and irregular halos at the spheroid rim; while this irregularity was only qualitatively observable in the 2D brightfield projection, sphericity computed from the full 3D segmented OCT volume offers a quantitative, volumetric counterpart to this rim irregularity. The two gray-level non-uniformity texture features, from the intensity and variance volumes respectively, likely capture complementary structural and dynamic heterogeneity across the spheroid volume, while SV-derived first-order skewness indicates shifts in the distribution of dynamic contrast within the spheroid, consistent with spatially heterogeneous response patterns.

The repeat-acquisition robustness test supports that the extracted features are not dominated by acquisition noise or unstable processing. Across six repeated scans of the same spheroid, relative dispersion (IQR/median) was low and comparable across all four selected features (0.032 to 0.047; Table 1), indicating stable extraction under unchanged conditions for both the shape/first-order descriptors and the gray-level non-uniformity texture features. This combination of low repeat-scan dispersion and substantially larger cohort-level variability (Table 1) indicates that the spread observed across the cohort reflects genuine between-spheroid differences rather than measurement noise, supporting the discriminative potential of these features for dose-dependent spheroid response.

The OCT+BF versus OCT-only comparison reached statistical significance only for Random Forest, and even there with comparatively low power (0.62); for Extra Trees, SVR, XGBoost, and LightGBM, the difference was not statistically significant and for SVR and XGBoost the OCT-only cross-validation IQR was in fact narrower than that of the corresponding OCT+BF model, indicating more consistent fold-to-fold performance for OCT-only. OCT radiomics alone already captures most of the dose-related signal present in the combined feature set. From a practical perspective this favors an OCT-only pipeline for dose-correlation modeling of spheroid radiation response, since it achieves comparable, and in some cases statistically indistinguishable or numerically superior, performance without requiring concurrent brightfield acquisition or manual annotation. The asymmetry discussed above, namely that brightfield features encode an explicit pre-irradiation baseline while the OCT acquisition in this study does not, may still explain the small residual advantage observed for Random Forest, but is evidently insufficient to produce a consistent, practically meaningfu benefit.

Several limitations should be considered. First, the study used a cross-sectional design (different spheroids per time point), which avoids repeated handling but prevents true within-spheroid longitudinal modeling and increases sensitivity to inter-spheroid variability. Second, the dataset is restricted to one cell line and a single experimental setup; generalization to other spheroid models and imaging systems remains to be tested. Third, dose correlation is a convenient standardized target but remains an indirect surrogate for biological endpoints; linking radiomic signatures to mechanistic readouts (e.g., viability, proliferation, hypoxia markers) would strengthen interpretability. Finally, segmentation quality and OCT acquisition parameters can influence radiomics; while curated masks reduce obvious artifacts, fully automated, standardized preprocessing is important for reproducibility at scale. Relatedly, the voxel size used for OCT acquisition (6.0 µm lateral, 3.5 µm axial in this study) determines the biological scale captured by each voxel: at this resolution, a single voxel can encompass multiple cells, such that texture features reflect aggregate, multi-cellular intensity and variance patterns rather than single-cell properties. A coarser or finer voxel size would sample a different underlying spatial scale, ranging from sub-cellular to multi-cellular, and could therefore yield a different and not directly comparable set of informative radiomic features. This scale-dependence should be considered when comparing radiomic signatures across OCT systems or acquisition settings, and motivates explicit reporting of voxel size, and ideally its harmonization, in future OCT radiomics studies. The choice of cell line is an additional factor that interacts with this scale-dependence, since cell size and morphology vary across cell lines, the same voxel size can correspond to a different number of cells per voxel, and consequently a different texture interpretation, depending on the cell line under study. Beyond this scale-related effect, different cell lines can also differ intrinsically in radiosensitivity, growth kinetics, and microenvironmental dynamics, such that the radiomic signatures and dose-response relationships identified here for SAS spheroids may not directly transfer to other cell lines, independent of any voxel-size considerations. Rather than viewing this purely as a limitation, this scale-dependence also suggests an opportunity: acquiring or resampling OCT volumes at multiple voxel sizes could provide complementary, scale-specific radiomic information, ranging from sub-cellular to multi-cellular descriptors, that a single fixed resolution cannot capture on its own, analogous to multi-scale or multi-resolution feature extraction approaches used in other radiomics domains Future work could explore whether combining radiomic features across multiple acquisition or resampling scales improves dose-response characterization beyond what is achievable at a single resolution.

In summary, OCT-derived radiomics provided a stronger and more consistent basis for dose prediction than brightfield morphology alone, and a small set of shared predictors emerged across model families that combined macroscopic size dynamics with OCT-based shape, texture, and SV-derived heterogeneity information. The repeat-scan experiment further indicated that these key descriptors are reproducibly extracted under unchanged acquisition conditions and that cohort-level variability substantially exceeded repeat-scan variability, supporting that the pipeline captures genuine, dose-related differences rather than predominantly measurement noise. Future work should prioritize (i) enabling true longitudinal OCT (e.g., reduced handling or incubator-coupled imaging) to track within-spheroid trajectories, (ii) validating the pipeline across additional cell lines, seeding regimes, and independent experimental batches, and (iii) linking radiomic signatures to orthogonal biological endpoints to strengthen interpretability. Methodological extensions such as automated segmentation with standardized quality assurance and time-series modeling of multi-day feature trajectories may further improve robustness and sensitivity to subtle treatment effects. Overall, OCT radiomics represents a sensitive, reproducible, and label-free high-throughput readout of spheroid radiation dose response that complements, and in many cases surpasses, conventional brightfield morphology

## Declarations

## Supplementary information

Not applicable.

## Ethics approval and consent to participate

Not applicable. This study used the established SAS human tongue squamous cell carcinoma cell line and did not involve human participants, identifiable human material, or animals.

## Consent for publication

Not applicable.

## Availability of data and materials

The datasets generated and analysed during the current study are available from the corresponding author on reasonable request.

## Authors’ contributions

MA designed the analysis pipeline, performed the imaging experiments and data analysis, and drafted the manuscript. RH and US contributed to the cell culture and spheroid experiments. LT contributed to the radiobiology methodology and interpretation. AT contributed to the OCT data processing. JS contributed to the radiation-physics methodology and interpretation. WN and MFS supervised the study and contributed to the imaging and radiomics methodology. All authors read and approved the final manuscript.

## Acknowledgements

We thank Dr. Ina Kurth (German Cancer Research Center Heidelberg, Germany) for providing the SAS human tongue squamous cell carcinoma cell line and for granting access to the Faxitron MultiRad 225 irradiator. The authors made limited use of large language model (LLM)–based AI tools for language editing of the manuscript. These tools did not generate any scientific results. The authors reviewed all output and take full responsibility for the final content.

